# AAV-mediated peripheral single chain variable fragments’ administration to reduce cerebral tau in adult P301S transgenic mice: mono- vs combination therapy

**DOI:** 10.1101/2025.02.13.638144

**Authors:** Sebica Katel, Julia Cicalo, Valeria Vasciaveo, Joseph Carrion, Mahadeo Leann, Patricio T. Huerta, Philippe Marambaud, Luca Giliberto, Cristina d’Abramo

**Affiliations:** The Litwin-Zucker Center for Alzheimer’s Disease & Memory Disorders, Institute of Molecular Medicine, Feinstein Institutes for Medical Research, Manhasset, NY, USA; Laboratory for Behavioral and Molecular Neuroimaging (LBMN) Center for Neurosciences, Feinstein Institutes for Medical Research, Manhasset, NY, USA; Laboratory of Immune & Neural Networks, Feinstein Institutes for Medical Research, Manhasset, NY, USA; Elmezzi Graduate School of Molecular Medicine at Northwell Health, Manhasset, NY, USA; Department of Molecular Medicine, Zucker School of Medicine at Hofstra/Northwell, Hempstead, NY, USA; Donald and Barbara Zucker School of Medicine at Hofstra/Northwell, Hempstead, NY, USA; Institute for Neurology and Neurosurgery, Northwell Health, Manhasset, NY, USA

**Keywords:** tau, immunotherapy, scFv, gene therapy, monotherapy, combination therapy Alzheimer’s disease

## Abstract

Tau is a primary target for immunotherapy in Alzheimer’s disease. Recent studies have shown the potential of anti-tau fragment antibodies in lowering pathological tau levels *in vitro* and *in vivo*. Here, we compared the effects of single-chain variable fragments (scFv) derived from the well-characterized monoclonal antibodies PHF1 and MC1. We used adeno-associated virus 1 (AAV1) to deliver scFvs to skeletal muscle cells in eight-week-old P301S tau transgenic mice. We evaluated motor and behavioral functions at 16 and 23 weeks of age and measured misfolded, soluble, oligomeric and insoluble brain tau species. Monotherapy with scFv-MC1 improved motor and behavioral functions more effectively than scFv-PHF1 or combination therapy. Brain glucose metabolism also benefited from scFv-MC1 treatment. Surprisingly, combining scFvs targeting early (MC1) and late (PHF1) tau modifications did not produce additive or synergistic effects. These results confirm that intramuscular AAV1-mediated scFv-MC1 gene therapy holds promise as a potential treatment for Alzheimer’s disease. Our findings also suggest that combining scFvs targeting different tau epitopes may not necessarily enhance efficacy if administered together in a prevention paradigm. Further research is needed to explore whether other antibodies’ combinations and/or administration schedules could improve the efficacy of scFv-MC1 alone.

**Graphical abstract/eTOC synopsis:** Katel and colleagues show that peripheral vectorized scFvMC1 (in monotherapy) reduces pathological tau species in tau transgenic mice more efficiently than in combination with scFv-PHF1. The authors observed improved motor and behavioral functions together with increased brain glucose metabolism in scFv-MC1-treated mice.

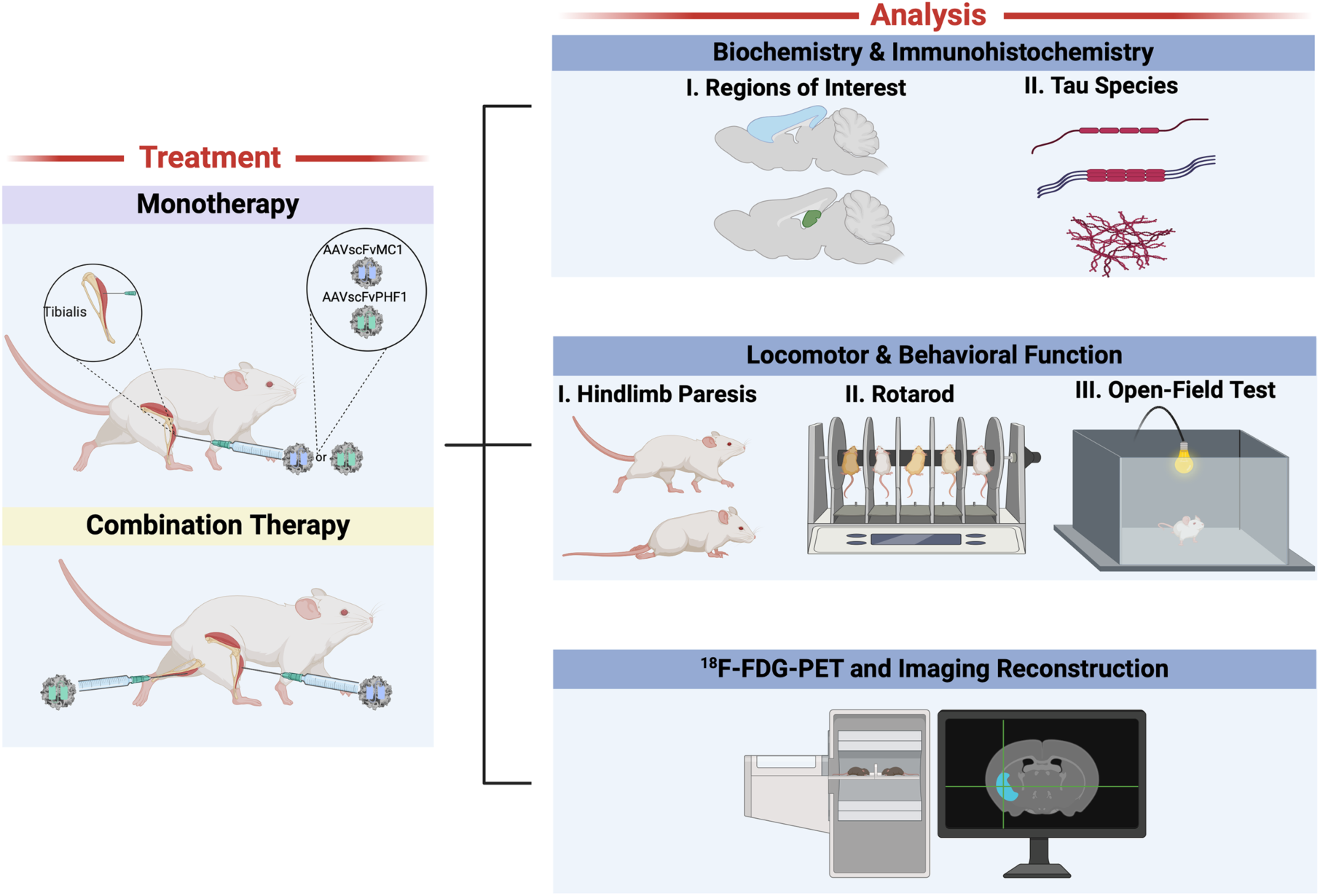

## Introduction

In Alzheimer’s disease (AD), the development of disease-modifying therapies targeting MAPT (microtubule-associated protein tau) is supported by the strong correlation between tau deposition and the onset and progression of cognitive decline (1–5). While not present in AD, MAPT gene mutations cause tau accumulation in hyperphosphorylated, conformationally altered, and aggregated form, leading to neurodegeneration, e.g. in frontotemporal dementia (6–8). Reducing pathological tau is a promising strategy, and a wide array of approaches have been proposed and tested, aiming to clear tau aggregates, slowing the progression of the disease: immunotherapy, inhibition of aggregation, autophagy stimulation, modulation of phosphorylation, glycosylation, acetylation, truncation and antisense oligonucleotides (ASO) (9–13). While Aβ immunotherapy has progressed towards regulatory approval (14, 15), with modest benefits and potential side effects, tau therapeutics are still in the early phases of clinical development (9, 16).

Current anti-tau immunotherapy use whole IgGs, which poorly penetrate the blood-brain barrier (17, 18). Anti-tau antibodies targeting extracellularly the N-terminus of tau, have failed to provide benefits in human trials (9, 11, 19, 20), raising two critical questions: 1) are these antibodies targeting relevant epitopes of tau, at the right time? and 2) is focusing solely on extracellular tau sufficient for therapeutic effects (21)? The recent phase II trial of Zagotenemab (NCT03518073), a humanized form of the MC1 antibody (conformational-tau specific, early marker of AD pathology) (22, 23), did not meet its primary endpoint and was discontinued (9). This result was attributed to the antibody’s poor binding to intracellular tau (21). It is important to note that preclinical studies in tauopathy models have demonstrated some ability of antibodies to enter neurons (24–26).

A new avenue is represented by antibody fragments, which, by virtue of their small size, readily enter the brain, diffuse into the parenchyma, and bind to complex epitopes, often unrecognized by whole antibodies (27–29). Engineered anti-tau antibodies employed as *in vivo* diagnostic tools have confirmed neuronal uptake and intracellular distribution after peripheral administration (28, 30, 31), with signals clearly correlating with intraneuronal tau aggregates, associated with markers of endosomes, autophagosomes, and lysosomes. Using antibody fragments, such as single-chain variable fragments (scFv), intrabodies (IBs), and variable domains of heavy-chain antibodies (VHH), has shown great potential as disease-modifying therapeutics *in vitro* and *in vivo* (28, 29, 32–42). Antibody engineering combined with vectorized gene therapy represents an important alternative approach to delivering anti-tau fragments to the brain parenchyma. Our laboratory has shown that AAV-vectorized scFv-MC1 can significantly reduce pathological tau species in adult tau transgenic mice upon AAV-mediated hippocampal injection (32). In addition, we have optimized a translational preclinical paradigm in which a one-time intramuscular injection of AAV1-scFv-MC1 was able to generate a long-lasting peripheral source of scFv-MC1, translating into a parallel reduction of tau species in the brain (33) with an overall efficacy between 50% and 60%.

Here, we sought to validate further and enhance the efficacy of this paradigm, providing a comparison between scFv-MC1, scFv-PHF1, and their combination. The use of PHF1 (phosphorylated tau-specific, p-Ser396/404, late epitope), has proven effective in reducing pathological tau in transgenic models (43, 44). In addition, intracranial injection in adult P301S mice using AAVrh.10-PHF1 (carrying the full-length mAb) has shown robust (>70%) reductions of insoluble phosphorylated tau (p-tau) species in the hippocampus and cortex (35). This study explores whether a combination approach targeting both AD specific early-conformational modifications (scFv-MC1) and late phosphorylation epitopes (scFv-PHF1) would result in additive efficacy. A detailed biochemical analysis of misfolded, soluble and insoluble tau species in cortex and hippocampus shows that scFv-MC1 outperforms all treatments paradigms, with improved motor and behavioral phenotypes. We also observed that scFvMC1 treatment is associated with an increased relative glucose metabolic uptake in specific brain areas relevant to AD pathology. These new data extend our knowledge on tau immunotherapy with special emphasis on 1) which tau biochemical species are targeted or engaged and their correlation with the neurodegenerative phenotype and 2) which (if any) combination approach will enhance efficacy when targeting tau.

## Results

### scFv-PHF1 retains its epitope specificity and ability to bind pathological tau

While scFv-MC1 was fully characterized previously (32, 33), in this study, we assessed the ability of scFv-PHF1 to target phosphorylated tau at the 396/404 epitope. After 48h of transfection into HEK293T, scFv-PHF1 was efficiently secreted into the conditioned medium as shown by its reactivity to the specific p396/404 epitope (Figure 1Aa). Cell lysates from transfected HEK293T cells revealed the scFv by immunoblot (with an expected band around 30 kDa) (Figure 1Ab). Furthermore, purified scFv-PHF1 was analyzed on a Coomassie-stained SDS-PAGE gel to confirm its molecular weight and the purity of the preparation (Figure 1Ac). To further characterize scFv-PHF1, we stained brain sections from 5-month-old P301S tau transgenic mice using either PHF1 (full IgG1) or the purified scFv-PHF1 (Figure 1B), confirming its ability to bind tangle-like bearing neurons efficiently. Additionally, reactivity to aggregated/phosphorylated tau was measured using an immunosorbent assay in which 96-well plates were coated with AD-derived PHF-tau (Figure 1C). As expected, scFv-PHF1 showed greater reactivity to PHF-tau than scFv-MC1, confirming previous affinity data published on the parental mAbs (22). Collectively, these results demonstrate that scFv-PHF1 retains its epitope specificity and ability to bind pathological tau, suggesting its potential as a disease-modifying therapeutic.

**Figure 1.**
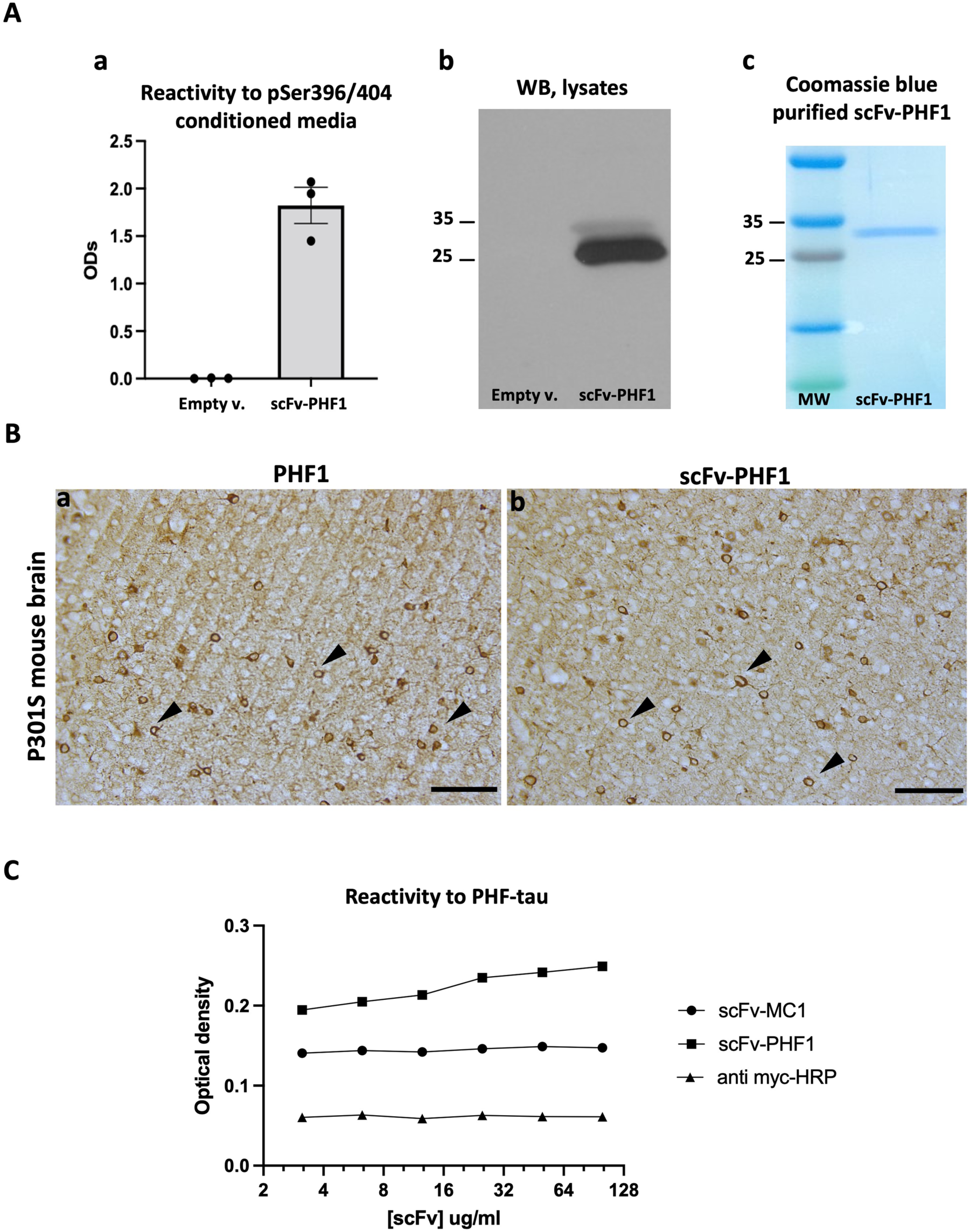
ScFv PHF1 characterization of epitope specificity and target engagement. **(A) a)** scFv-PHF1 was successfully secreted upon transfection in HEK293T cells, shown by strong reactivity of the conditioned medium to the p396/p404-tau peptide; no signal was detected in the empty vector lane (Empty v). **b)** Cell lysates were tested via immunoblot for scFv detection at 30 kDa (anti-Myc tag primary antibody); no band was detected in the empty vector lane (Empty v). **c)** Purified scFv-MC1 was run on a Coomassie-stained SDS page confirming its molecular weight (30 kDa). **(B)** a) PHF1 and b) scFv-PHF1 stain tangle-bearing neurons in P301S tau transgenic mice brains; staining was visualized using 3,3’-Diaminobenzidine (DAB); (Olympus BH-2 bright field microscope, bar: 100 μm). **(C)** Specificity of scFv-PHF1 was confirmed via immunosorbent assay coated with AD-derived PHF-tau; increasing concentrations (μg/ml) of scFvs (scFvPHF1 and scFMC1) were added following incubation with anti myc-HRP: reactivity to PHF-tau is expressed as Optical density; anti myc-HRP show the basal background in the assay.

### Expression of scFv-PHF1 and scFv-MC1 in monotherapy and combination significantly reduces both misfolded and soluble tau in the cortex

To investigate the effect of the different treatment paradigms, we initially focused on the levels of MC1-misfolded tau in the cortex. Immunohistochemical analysis and semi-quantification of tau pathology using MC1 mAb were employed for this purpose. Our data showed that all treatment groups significantly reduced the accumulation of misfolded tau in the frontal cortex (Figure 2 A, B): a 45-50% decrease was detected in the scFv-PHF1 injected group (* *P* < 0.05), a 60% reduction in the scFvMC1 group (*** *P* < 0.001) and 45-50% in combination (** *P* < 0.01). To deepen our analysis, we rigorously quantified the levels of total and phosphorylated soluble tau using our highly sensitive tau ELISAs (45). As shown in Figure 2C, a conspicuous decrease of total and p-tau was detected in the cortex. When compared to control mice (Sham eGFP), total soluble tau was reduced by circa 60% in the scFvPHF1 and scFvMC1 groups (**** *P* < 0.0001), while their combination exerted a 70% reduction (**** *P* < 0.0001). Animals treated in combination also showed significantly less total tau compared to the scFv-PHF1 group (* *P* < 0.05). We then looked at phosphorylated tau in the same conditions and found that tau phosphorylation at Ser202 and Ser396/404 followed the same trend as total tau, with both phospho-epitopes decreased by 60% (*** *P* < 0.001, **** *P* < 0.0001) in all treatment groups. Conversely, phosphorylation at Thr231 was not significantly modulated. Of note, while the levels of total tau and pSer202 were comparable in the baseline (2-month-old mice, not injected, termed ‘2M’ henceforth) and sham groups (6-month-old mice, not injected), pThr231 and pSer396/404 showed an expected boost of phosphorylation with aging (2 to 6 months). Examining the data from this perspective, all treatment paradigms were able to reduce total and pSer202 tau to significantly lower levels than the 2M baseline group (−50% to −70%), implying the ability to actively remove existing soluble tau species. In addition, the buildup of tau phosphorylated at Ser396/404 was contained by all treatments, as with levels of p-tau comparable to the 2M baseline group. Taken together these data show that, in the cortex, all treatment groups were able to efficiently target MC1-misfolded, total, and phosphorylated soluble tau, both clearing up and blocking the build-up of newly phosphorylated species. No sex differences were detected in this analysis (Supplementary Figure 1).

**Figure 2.**
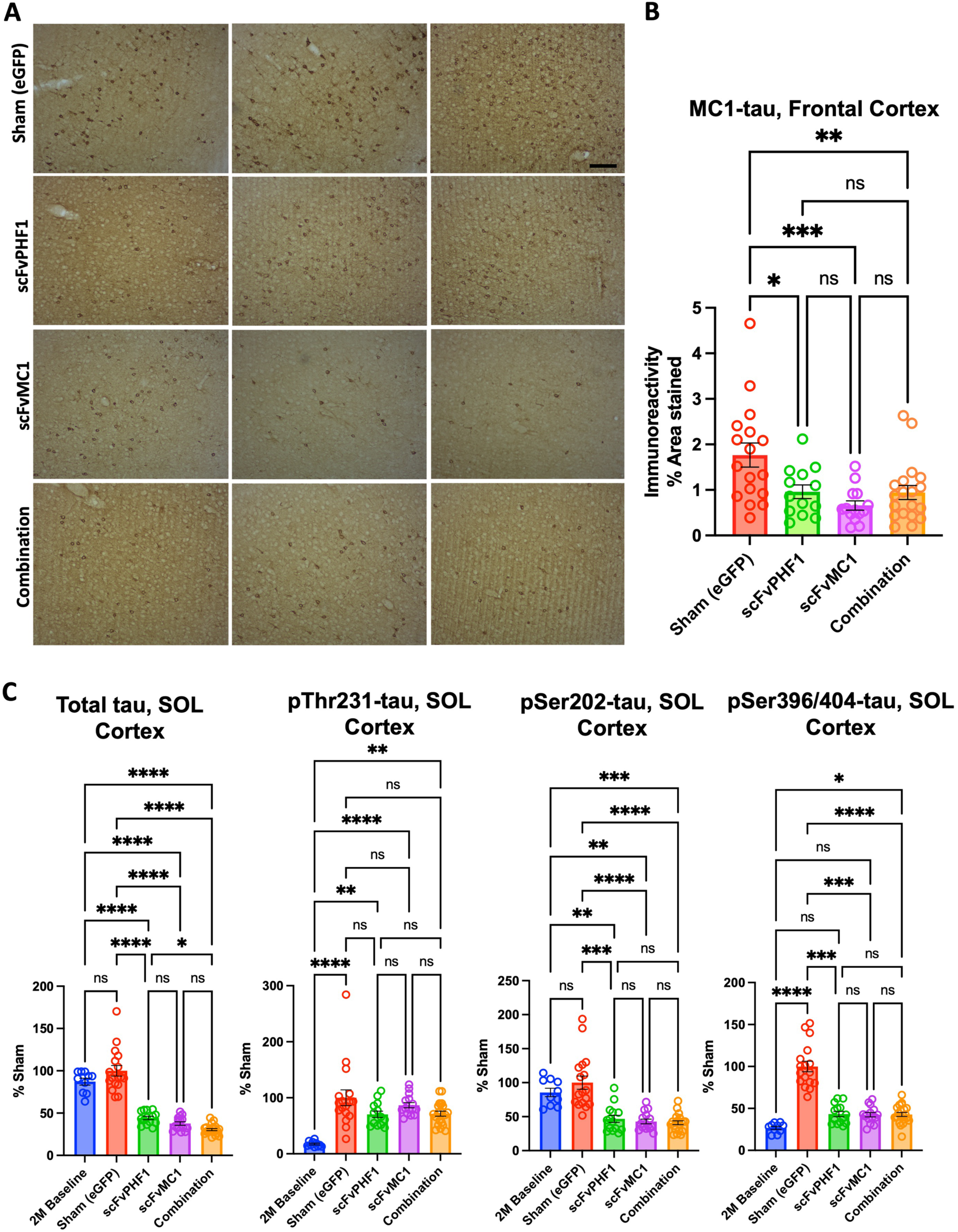
scFv-MC1 and scFv-PHF1 significantly reduce misfolded and soluble tau in the cortex, in mono- and combination therapy. **(A)** Representative images of MC1-misfolded tau in frontal cortex (Olympus BH-2 microscope, bar: 100 μm), and **(B)** semiquantification (% area stained). Treatment groups: sham eGFP (n = 17), scFv-PHF1 (n = 13), scFv-MC1 (n = 14), and combination (n = 20). **(C)** ELISA quantifications of soluble (SOL) total tau, pThr231-tau, pSer202-tau, and pSer396/404-tau in 2-month-old baseline (n = 10), sham-eGFP (n = 17), scFv-PHF1 (n = 14), scFv-MC1 (n = 15), and combination (n = 21). Data were normalized to the percent sham-eGFP group and represent mean ± SEM. Statistical analysis was performed by non parametric Mann-Whitney test. * *P* < 0.05; ** *P* < 0.01; *** *P* < 0.001; **** *P* < 0.0001.

### Misfolded tau, but not soluble tau, is decreased in the hippocampus when expressing scFv-MC1 in monotherapy

Semi-quantification of tau pathology using the MC1 mAb in the hippocampus showed that the granule cell layer of the dentate gyrus (Figure 3 A, B) exhibited a 70% reduction of reactivity in the group receiving scFv-MC1 monotherapy compared to the sham group (* *P* < 0.05). Conversely, scFv-PHF1 and its combination with scFvMC1 delivered a nonsignificant trend toward reduction. In a parallel biochemical analysis of the whole hippocampus, total soluble tau was not modulated when compared to the sham injected mice, except for a minimal trend to reduction in the scFvMC1 and combination cohorts (Figure 3C); however, scFvMC1 and combination-treated mice had significantly less total tau than the scFvPHF1 injected group (* *P* < 0.05). The animals from the 2M baseline cohort showed overall the same amount of total tau as the sham mice but significantly less tau phosphorylation at Ser202 and Ser396/404. None of the treatment paradigms were able to block or tamper down the buildup of phosphorylated soluble tau in the hippocampus. Similar to the cortex, no sex effect was found in the hippocampus (Supplementary Figure 2). These data indicate that, in the hippocampus, while the overall amount of total and phosphorylated soluble tau were not efficiently modified by any of the treatment options, MC1-misfolded tau was substantially reduced in the dentate granule cells, a hippocampal subregion implicated in learning, memory, and spatial navigation (46, 47).

**Figure 3:**
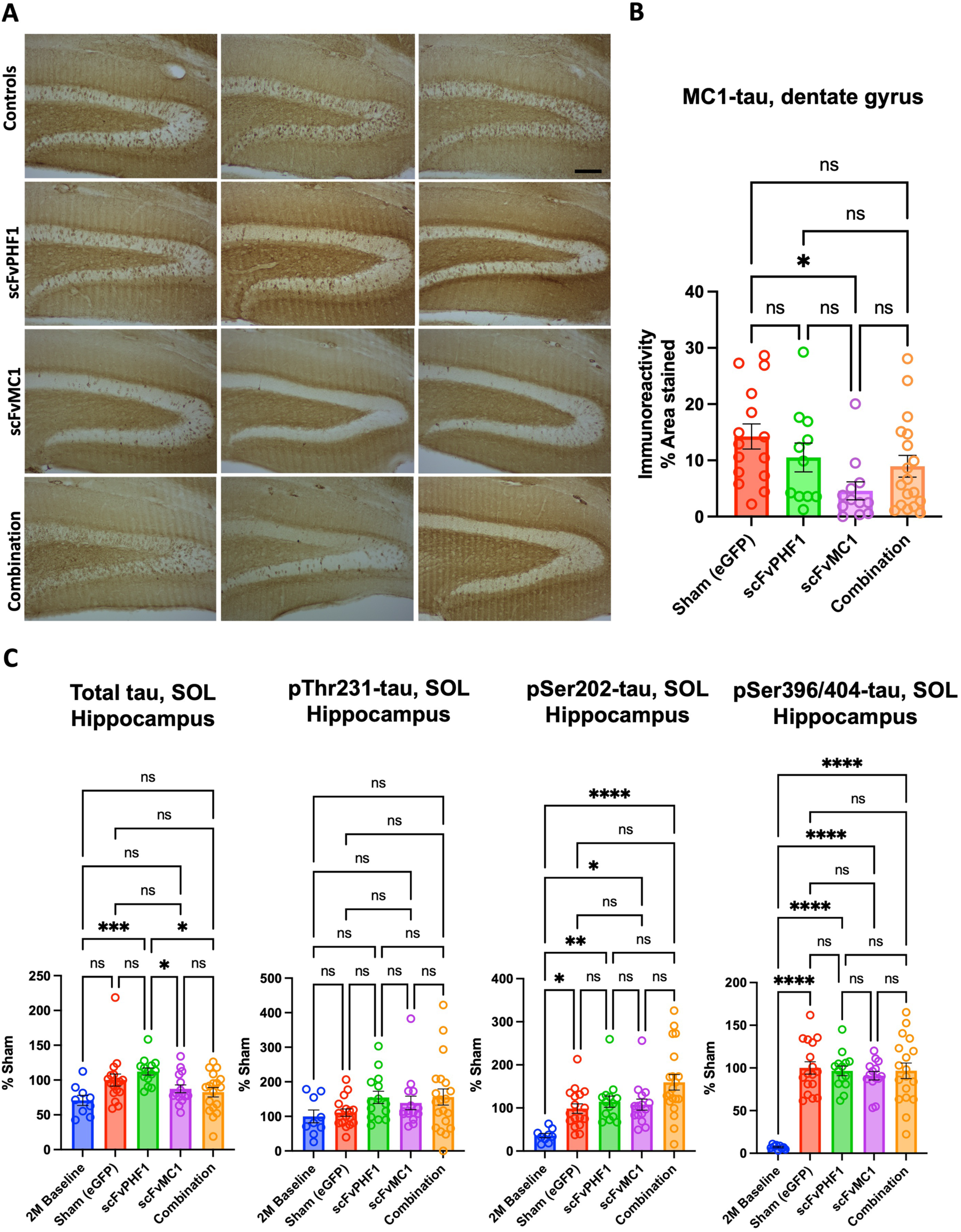
scFv-MC1 in monotherapy reduces misfolded tau in the hippocampus, while soluble tau levels remain stable. **(A)** Representative images of MC1-misfolded tau in dentate gyrus (Olympus BH-2 microscope, bar: 100 μm). (**B)** Semi-quantification of percent area stained per treatment group: sham-eGFP (n = 16), scFv-PHF1 (n = 13), scFv-MC1 (n = 14), and combination (n = 20). **(C)** ELISA quantifications of soluble (SOL) total tau, pThr231-tau, pSer202-tau, and pSer396/404-tau in 2-month-old baseline (n = 9), sham-eGFP (n = 17), scFv-PHF1 (n = 14), scFv-MC1 (n = 15), and combination (n = 18). Data were normalized to the percent sham-eGFP group and represent mean ± SEM. Statistical analysis was performed by one-way ANOVA or non parametric Kruskal-Wallis test. * *P* < 0.05; ** *P* < 0.01; *** *P* < 0.001; **** *P* < 0.0001.

### scFv-MC1 outperforms all treatment paradigms by significantly decreasing oligomeric/aggregated tau in both cortex and hippocampus

We next focused on oligomeric/aggregated tau species ranging from dimers to larger aggregates. In Figure 4A,B (females and males combined, F&M) oligomeric tau was significantly lowered upon scFv-MC1 expression in cortex (40% reduction, ** *P* < 0.01) and hippocampus (50% reduction, **** *P* < 0.0001) when compared to the sham group; while scFvPHF1 monotherapy exerted a non-significant trend toward reduction in cortex, it significantly reduced tau in the hippocampus (∼30%, ** *P* < 0.01); combining the scFvs decreased oligomeric tau in the hippocampus (30–35%, ** *P* < 0.01) with no effect in cortex. When breaking up the data by sex (Figure 4A), scFv-MC1 treatment lost its significance in cortex, yet preserved a trend to reduction, likely in relation to a reduced sample size. Notably, scFv-MC1 mantained its efficacy in the hippocampus (Figure 4B) of both females and males, reaching −60% (*** *P* < 0.001) and −40% (** *P* < 0.01) reduction, respectively. Once again, no significant sex differences were evident. Altogether, these data confirmed that scFv-MC1, in monotherapy, significantly and consistently engaged oligomeric/aggregated tau in the hippocampus and cortex, and that no additive effect resulted from the combination of the two scFvs. In addition, oligomeric tau in the hippocampus reached levels comparable to the 2M baseline group when expressing scFv-MC1, confirming the ability of this therapeutic tool to actively target and block the formation of aggregated species.

**Figure 4.**
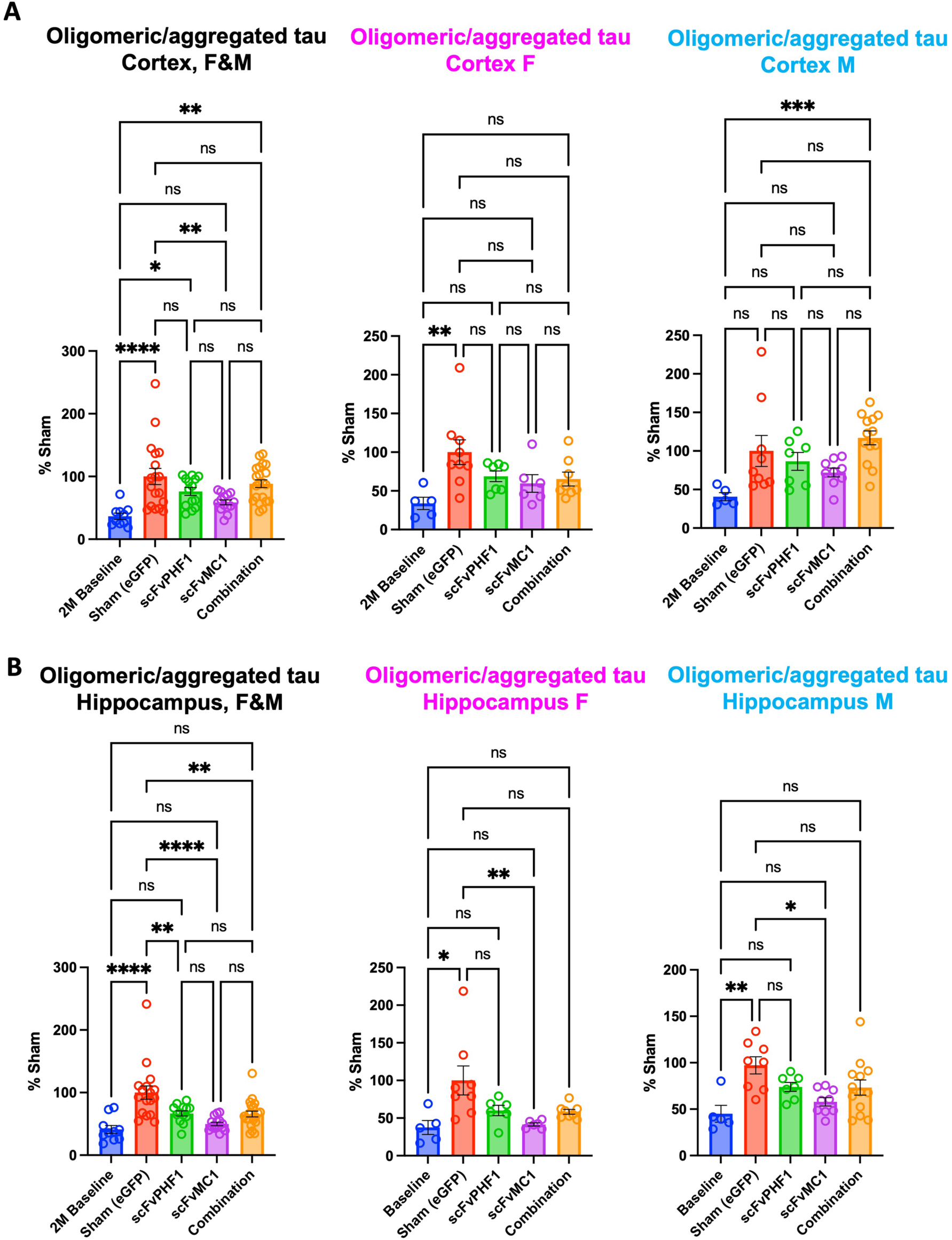
Cortical and hippocampal oligomeric/aggregated tau is significantly reduced with the scFv-MC1 treatment paradigm. **(A)** DA9/DA9-hrp sensitive tau ELISA quantified cortical oligomeric tau of combined (female [F] and male [M]) and separated sexes. Treatment groups: baseline (n = 10, 5F and 5M), sham-eGFP (n = 18, 9F and 9M), scFv-PHF1 (n = 14, 7F and 7M), scFv-MC1 (n = 15, 6F and 9M), and combination (n = 21, 8F and 13M). **(B)** Hippocampal oligomeric tau quantifications of combined (F and M) and separated sexes. Treatment groups: baseline (n = 10, 5F and 5M), sham-eGFP (n = 16, 8F and 8M), scFv-PHF1 (n = 13, 6F and 7M), scFv-MC1 (n = 15, 6F and 9M), and combination (n = 21, 8F and 13M). Data were normalized to the percent sham-eGFP group and represents mean ± SEM. Statistical analysis was performed by one-way ANOVA or non parametric Kruskal-Wallis test. * *P* < 0.05; ** *P* < 0.01; *** *P* < 0.001; **** *P* < 0.0001.

### scFv-MC1 reduces insoluble tau in the hippocampus but not in the cortex

None of the treatments were able to decrease insoluble tau in the cortex (Figure 5A), despite the reduction of oligomeric/aggregated tau in the scFv-MC1 group (Figure 4A). However, combining the scFvs resulted in increased levels of total insoluble tau (>80%, **** *P* < 0.0001). We then sought to investigate whether insoluble tau was modulated in the hippocampus. Given that scFv-MC1 monotherapy proved to be the most effective treatment paradigm in this study, we treated a new cohort of mice with scFv-MC1 with the purpose of extracting insoluble tau from the hippocampi, with hippocampi from each mouse being pooled in order to obtain sufficient amounts of insoluble preparation. As shown in Figure 5B, total insoluble tau was reduced by 40% in the scFv-MC1 group compared to the sham-injected mice. In addition, the treated mice reached levels comparable to the 2-month baseline (*P* = 0.2502, ns). Phosphorylated tau showed a nonsignificant trend to reduction, in particular for the pSer202 epitope (*P* = 0.0524).

**Figure 5.**
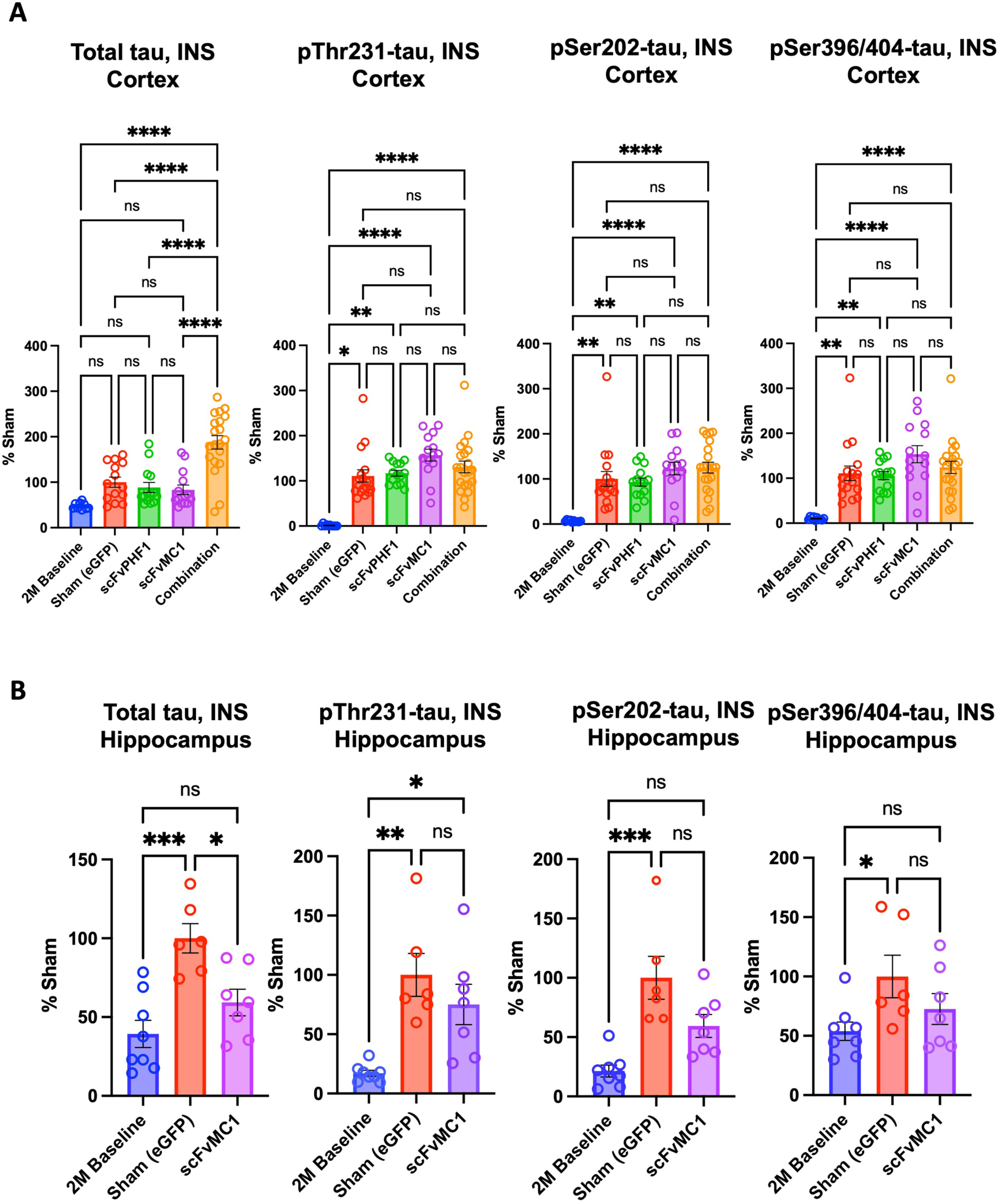
scFv-MC1 provides a reduction of insoluble tau in hippocampus, but fails to significantly reduce insoluble tau in cortex. **(A)** ELISA quantifications of cortical insoluble (INS) total tau, pThr231-tau, pSer202-tau and pSer396/404-tau in baseline (n = 10), sham-eGFP (n = 14), scFv-PHF1 (n = 14), scFv-MC1 (n = 14), and combination groups (n = 21). **(B)** ELISA quantifications of hippocampal insoluble (INS) total tau, pThr231-tau, pSer202-tau, and pSer393-404-tau in baseline (n = 8), sham-eGFP (n = 6), and scFv-PHF1 (n = 7). Data were normalized to the percent sham-eGFP group and represent mean ± SEM. Statistical analysis was performed by one-way ANOVA or non parametric Kruskal-Wallis test. * *P* < 0.05; ** *P* < 0.01; *** *P* < 0.001; **** *P* < 0.0001.

### Motor and behavioral phenotypes are ameliorated by scFv-MC1 in monotherapy

To assess the effect of disease modification, we examined mice for motor and behavioral functions. Given the early hind-limb paresis developing in the P301S model and shortened lifespan, mice were tested in a time frame where motor deficit would not affect their performance, allowing us to detect differences between groups. Mice (females and males) were studied at 16 weeks of age (8 weeks post-AAV-scFv injection) using the rotarod task to measure motor coordination, balance and grip strength. We found that the scFv-MC1 group showed significant amelioration of the motor phenotype when compared to the sham group (*** *P* = 2.05×10-^8^, 2-way RMANOVA with Tukey test), the scFv-PHF1 group (** *P* = 0.00128) and combination treatment (** *P* = 0.00124) (Figure 6A). In addition, animals treated with scFv-MC1 monotherapy had a significant lower degree of hind-limbs clasping compared to any other group at the time of sacrifice (26 weeks) (Figure 6B). The open field task examined locomotor activity and anxiety-like behavior at 16 weeks (trial 1, 8 weeks post-AAV-scFv injection) and 23 weeks (trial 2) (Figure 6C). This task was run only in males, to avoid any confounder due to poor locomotion in females who generally present significant clasping at 23 weeks of age. Overall, we found a significant rescue of the motor phenotype with scFv-MC1 treatment at trial 1 that persisted to trial 2, as shown by parameters such as distance moved (scFv-MC1 v. sham, *** *P* = 5.42×10^-6^; scFv-MC1 v. scFvPHF1, *** *P* = 1.67×10^-5^; scFv-MC1 v. combination, *** *P* = 0; 2-way RMANOVA with Tukey test), speed (scFv-MC1 v. sham, *** *P* = 4.49×10^-6^; scFv-MC1 v. scFvPHF1, *** *P* = 1.39×10^-5^; scFv-MC1 v. combination, *** *P* = 0), and mobility state (scFv-MC1 v. sham, *** *P* = 1.56×10^-5^; scFv-MC1 v. scFvPHF1, *** *P* = 2.03×10^-5^; scFv-MC1 v. combination, *** *P* = 0). Notably, the anxiety-like phenotypes were also rescued in the scFv-MC1 group which, compared to the other groups, spent significantly longer time in the center of the arena (scFv-MC1 v. sham, *** *P* = 3.16×10^-4^; scFv-MC1 v. scFvPHF1, *** *P* = 2.87×10^-6^; scFv-MC1 v. combination, *** *P* = 1.61×10^-6^) and significantly less time in the corner zones (scFv-MC1 v. sham, *** *P* = 5.18×10^-4^; scFv-MC1 v. scFvPHF1, *** *P* = 1.42×10^-5^; scFv-MC1 v. combination, *** *P* = 1.07×10^-4^). No significant benefit of scFv-MC1, scFv-PHF1 or their combination was found on cognitive measures, but we observed a trend toward a benefit with scFv-MC1 alone on the object-place memory task (data not shown) (48).

**Figure 6.**
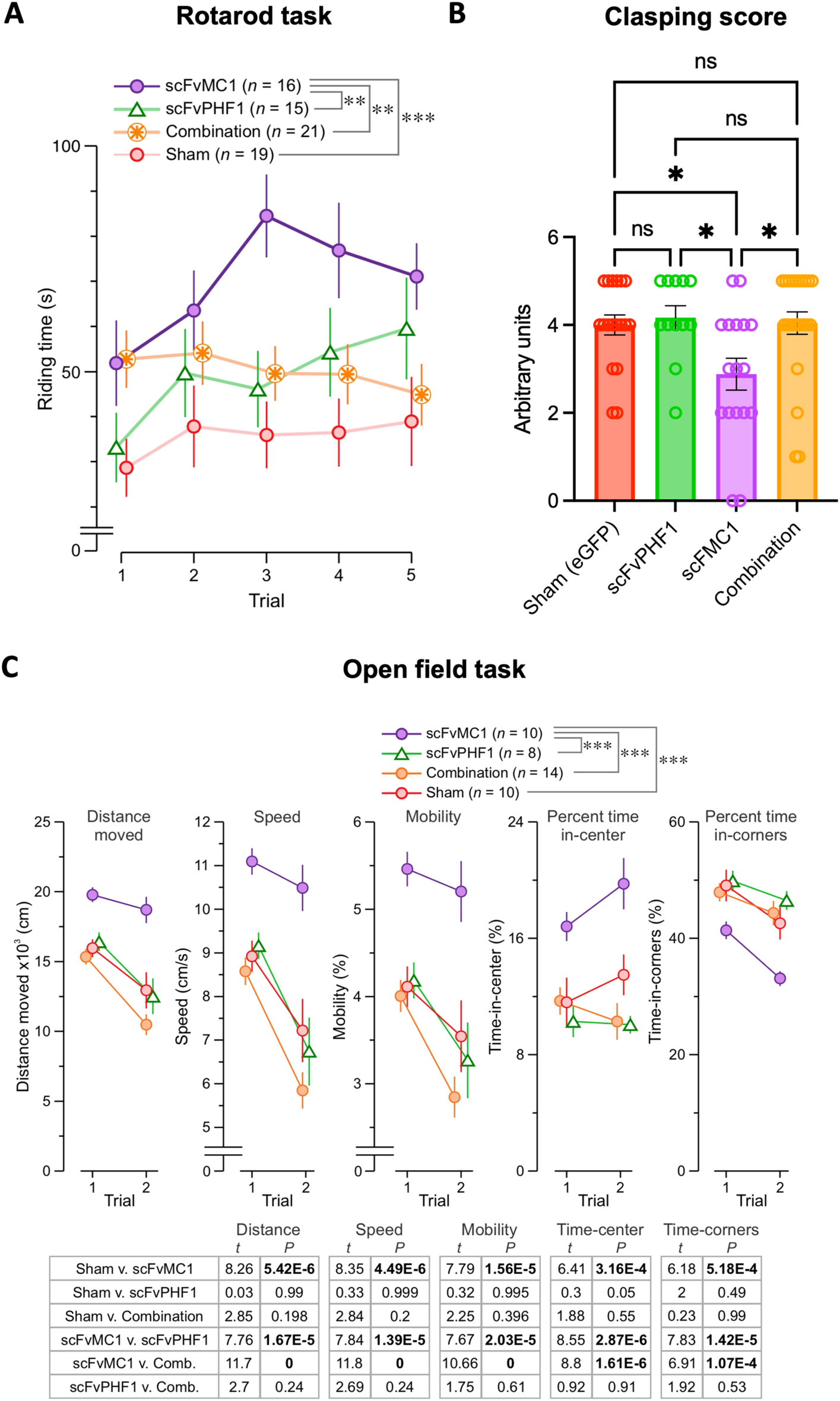
Motor and behavioral abnormalities are ameliorated by scFv-MC1 treatment. **(A)** Riding time in the rotarod task (5 trials) for 16-week-old mice (8 weeks post-AAV-scFv injection). Treatment groups: sham (n = 19, 9F and 10M), scFv-PHF1 (n = 15, 7F and 8M), scFv-MC1 (n = 16, 6F and 10M), and combination (n = 21, 9F and 12M). **(B)** Hind-limb clasping scale: score 0 = normal escape extention reflex; score 5 = absence of extension reflex in both hind-limbs; score 1 through 4 were determined on the basis of the amplitude of the extention and the number of limbs affected by clapsing. Treatment groups are the same as above. Data represent mean ± SEM. Statistical analysis was performed by one-way ANOVA or non parametric Kruskal-Wallis test. * *P* < 0.05. **(C)** The open field task assessed locomotor activity (total distance moved, mean speed, mobility state) and anxiety-like behavior (% time in center, % time in corner zones) in male mice at 16 weeks (trial 1, 8 weeks post-AAV-scFv injection) and 23 weeks (trial 2). Treatment groups: sham (n = 10), scFv-PHF1 (n = 8), scFv-MC1 (n = 10), and combination (n = 14). Data represent mean ± SEM. Statistical analysis was performed by one-way ANOVA with repeated measures. * *P* < 0.05; ** *P* < 0.01; *** *P* < 0.001.

### scFvMC1 treatment alters brain glucose metabolism

^18^F-FDG-PET reflects the rate of cerebral glucose metabolism which is mainly determined by synaptic activity (49). We first compared 5.5-month-old C57BL6 mice with aged matched P301S animals (Supplementary Figure 3), detecting reduced cerebral glucose metabolism in the lateral entorhinal cortex (** *P* < 0.01), presubiculum (** *P* < 0.01), piriform cortex (*P* = 0.0587, ns) and lateral striatum (** *P* < 0.01) in the tau transgenic mice (analysis run at *P* < 0.05), consistent with the neuronal pathology present in this animal model. Given the beneficial effect of scFvMC1 in this study, we then assessed whether scFvMC1 could affect glucose metabolic uptake in distinct brain regions, and treated a cohort of P301S mice with AAV1-scFvMC1 or AAV1-eGFP (Sham) (Figure 7 and Supplementary Figure 4). We found that the scFv treated mice exhibited an increased ^18^F-FDG uptake in the dentate gyrus (* *P* < 0.05), entopenducular nucleus (* *P* < 0.05; nonprimate homolog of the globus pallidus internus – GPi), reticular nucleus (* *P* < 0.05) and periaqueductal gray (* *P* < 0.05) when the analysis was run at *P* < 0.05, while the amygdala showed significant reduction (*** *P* < 0.001) in glucose metabolism when the analysis was run at *P* <0.01. The P-value and cluster cut-off threshold set spatial extent of activations to clusters containing greater than 50 - 100 voxels per cluster, which strengthens statistical power by reducing over-interpretation of data. Overall, our data indicate a general amelioration of glucose metabolism in scFv treated P301S compared to sham-treated mice in brain regions relevant to spatial memory consolidation (46), attention (50, 51), reward (52) and anxiety and fear response (53–55).

**Figure 7.**
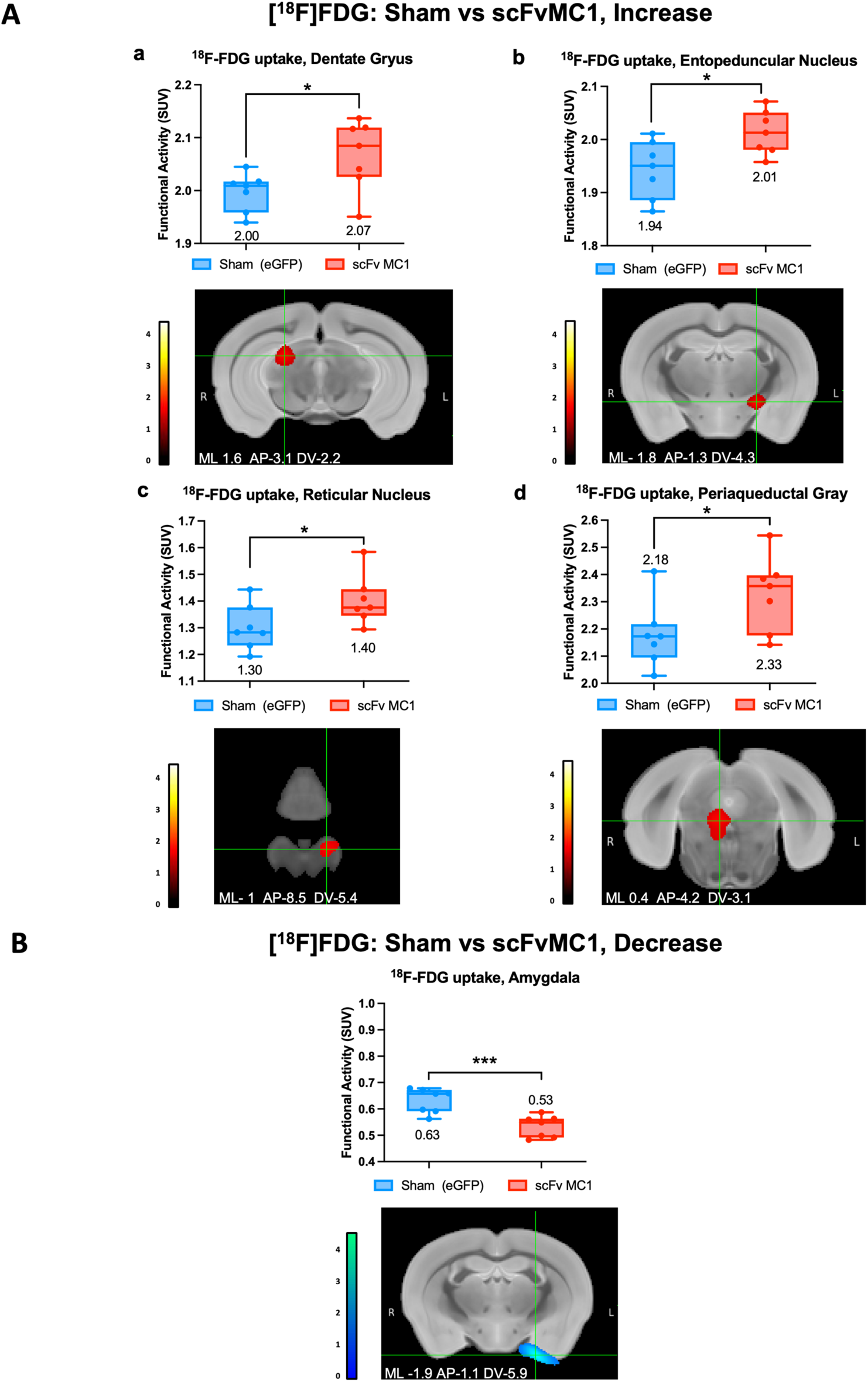
Regions with increased or decreased ^18^F-FDG uptake after scFvMC1 treatment. **(A)** Coronal view with mediolateral (ML), anterior posterior (AP), and dorsoventral (DV) coordinates shown with cluster overlay of statistically significant increase in ^18^F-FDG (*P* < 0.05) comparing scFvMC1 (P301S, n=7) vs. Sham mice (P301S, n=7). Brain regions with increased ^18^F-FDG uptake include: **(a)** dentate gyrus, **(b)** entopeduncular nucleus, **(c)** reticular nucleus, and **(d)** periaqueductal grey. **(B)** Coronal view of the amygdala with cluster overlay of statistically significant decrease (*P* < 0.01) in relative ^18^F-FDG uptake comparing scFvMC1 (P301s, n=7) vs. Sham mice (P301S, n=7). Significant clusters of activation from two-sample statistical comparison extracted using FSLeyes and overlaid onto coronal view mouse brain MRI template. Paxinos & Franklin anatomical coordinates shown with color bar representation of ^18^F-FDG increased or decreased uptake threshold value. Data are expressed as the mean + SEM. *p < 0.05, **p < 0.01, ***p < 0.001, two tailed unpaired t-test with Welch’s correction

## Discussion

The definitive pathological tau species responsible for neurodegeneration in AD and other tauopathies remain elusive. Consequently, it is also unclear whether targeting a single species or employing a combination approach would be more effective in anti-tau immunotherapies. In this *in vivo* study, we used a prevention protocol to compare the disease-modifying effect of AAV-mediated scFv-PHF1 and scFv-MC1 as monotherapy and in combination. To our knowledge, this is the first preclinical model exploring an anti-tau gene therapy combination approach. Our findings reveal that: 1) peripherally injected AAV-mediated scFv-PHF1 and scFv-MC1 equally reduce soluble tau in the cortex; 2) scFv-MC1 monotherapy is the only paradigm improving the motor and behavioral phenotype, demonstrating high translational relevance for tauopathies; 3) improved performance correlates with reduced oligomeric/aggregated and insoluble tau in the cortex and hippocampus, while cortical soluble tau reduction alone does not play a significant role; 4) scFvMC1 treatment increases glucose metabolism in brain areas involved in learning and memory consolidation; and 5) the lack of an additive or synergistic effects in the combination paradigm suggests that targeting both early and late modifications of tau is not necessarily beneficial, although other combinations of tau epitopes may enhance efficacy.

When using therapeutic antibodies directed to misfolded proteins like tau, it is crucial to understand how different aggregated species are modulated across brain regions. A regional analysis of tau aggregation states and post-translational modifications provides insight into how antibodies or their fragments act upon reaching the brain from the periphery. Figure 8A summarizes the effects of scFv-MC1, scFv-PHF1 and their combination on misfolded, soluble and oligomeric tau in the cortex and hippocampus, two regions relevant to AD. A striking observation is the difference in soluble tau modulation between the cortex and hippocampus. While all scFv-injected mice show significant reductions in cortical soluble tau, no changes are detected in the hippocampus. This effect is even more pronounced when comparing treated mice to the baseline cohort (Figure 2C): all therapeutic groups successfully cleared tau and/or blocked the buildup of newly phosphorylated species, reaching levels lower or equivalent to the 2-month-old baseline. Wenger *et al* (56) previously showed that P301S mice overexpress tau by 2.4-fold compared to healthy human controls, with soluble tau levels remaining steady in 2- and 5-month-old mice. Our data align with this observation, showing that baseline and sham groups have similar levels of total soluble tau. From this perspective, our strategy appears to exert a ‘reversion effect’ in lowering soluble tau in the cortex across all tested paradigms, suggesting a possible intracellular activity of the scFv.

**Figure 8.**
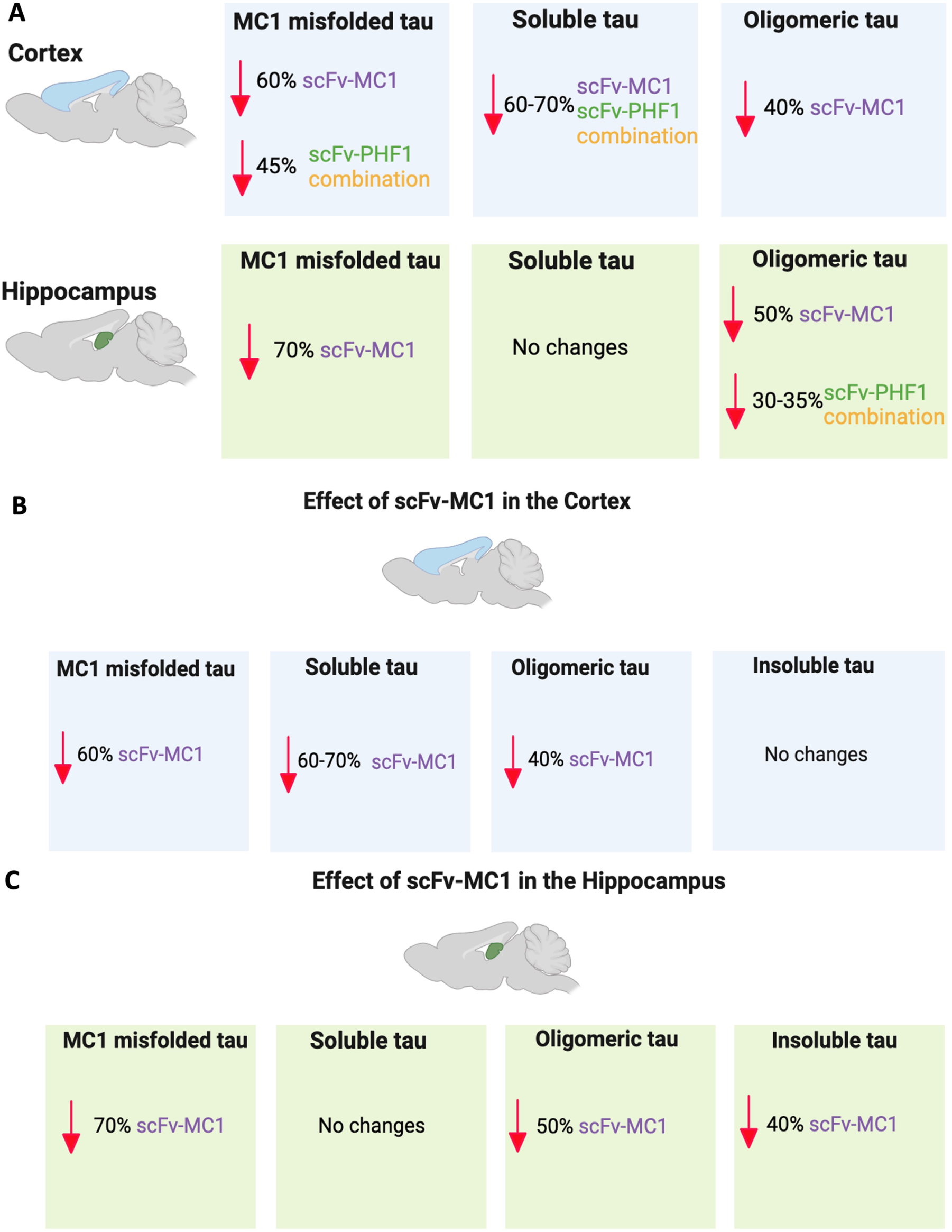
scFv-MC1 outperforms all other treatment paradigms in reducing MC1 misfolded, soluble, and oligomeric tau in AD relevant regions of cortex and hippocampus. **(A)** Overview of misfolded, soluble, and oligomeric tau post treatments of scFv-MC1, scFv-PHF1 and their combination. **(B)** MC1-misfolded, soluble, and oligomeric tau are all reduced in the cortex with scFv-MC1 treatment. **(C)** MC1-misfolded, oligomeric, and insoluble tau undergo reduction in the hippocampus with scFv-MC1 treatment.

The mechanism of action for peripherally injected scFvs targeting brain tau remains unclear. As in our previous study (33), we were unable to visualize scFvs in the brain (data not shown) due to detection sensitivity limits. We have previously shown that, upon intracranial injection of AAV, microglia are involved in clearing the scFv-tau immunocomplex, despite the lack of an Fc region (33). Additionally, a previous study (30) found that anti-tau scFvs co-localize with intracellular tau *in vivo* following peripheral injection. A recent article (38) reported that scFv-PHF1 might exert its action *in vitro* by entering cells and targeting intracellular tau, and we do not exclude the possibility that scFv-MC1 could act similarly. In our study, we hypothesize that the dramatic reduction in soluble tau in the cortical compartment results from scFvs entering neurons and acting on intracellular tau, as well as blocking extracellular species from ‘spreading’ between neurons. However, despite scFv-PHF1 and combination therapy significantly lowering soluble tau, this did not directly translate to improved behavioral phenotypes (Figure 6). Furthermore, the failure to reduce soluble tau in the hippocampus suggests that this species may be engaged according to a regional tropism. These observations raise questions about the relevance of decreasing overall soluble tau levels in tau immunotherapy, at least in mice, and highlight the importance of the quality and topography of targeted tau species over the total amount of tau.

To evaluate the potential clinical relevance of our study, we assessed mice for motor and behavioral performance and found that scFv-MC1 outperformed the other treatment paradigms (Figure 6), significantly improving motor and behavioral phenotypes compared to the other treatment groups. Goodwin *et al.* (38) have previously observed that scFv-PHF1 showed some degree of efficacy at delaying the onset of hindlimb paralysis. To our knowledge, this is the first evidence of a disease-modifying effect generated by an anti-tau scFv in tau transgenic mice, beyond delaying hindlimbs paralysis.

While it is difficult to establish a relationship between the topographical removal of tau in the regions described above, motor and behavioral changes and the FDG-changes, we do present a paradigm where scFv-MC1 treatment is able to alter and and normalize the metabolism of regions involved in memory, attention, fear, and reward. Several of these regions will not work in isolation, but rather in a network fashion (53): amygdala hyperactivation may be related to an anxiety phenotype (57); in our mice, the reduction of its activation, be it in a resting state, may underlie an amelioration of such network, and result in a normalization of their exploratory behavior.

Given its positive outcome on behavioral phenotypes and brain glucose metabolism, we focused our attention solely on the effect of scFv-MC1 in the cortex and hippocampus (Figure 8 B,C). We identified topographical differences in how the reduction of MC1-misfolded tau correlated with soluble or insoluble levels. In the cortex, the scFv appeared to target misfolded/soluble/oligomeric tau concurrently, with no impact on insoluble species (Figure 8B). In contrast, in the hippocampus, it was directed towards misfolded/oligomeric/insoluble species (Figure 8C), with no changes in the overall tau soluble levels. Thus, it is possible that MC1-misfolded tau is assembled differently in the cortex and hippocampus of P301S mice, leading to preferential accumulation of soluble or insoluble aggregates, respectively. This difference may result in distinct tropism of scFv-MC1 in different areas. To summarize this reasoning, scFv-MC1 might find a more readily available soluble pool of tau in the cortex (intracellular and extracellular) than in the hippocampus, resulting in an overall reduction of soluble tau. In the hippocampus, insoluble aggregates of MC1-misfolded tau might be more prevalent and accessible to the peripherally injected recombinant antibody, resulting in lower levels of the insoluble tau pool. We initially hypothesized that combining antibody fragments targeting early-conformational modifications and late-phosphorylation epitopes of tau could boost the efficacy of our system. However, our current data do not support this assumption. Despite the dual therapy’s ability to lower MC1-misfolded and soluble tau in the cortex to a considerable extent, this paradigm surprisingly led to an increase in total levels of insoluble tau (>80%) in the cortex. We do not know the reason for this effect. One could speculate that an excess of scFv binding to insoluble aggregates or their dimerization results in “stabilization” of the lesions, leading to an overall increase in insoluble levels.

To conclude, the present study demonstrates that AAV1-mediated scFv-MC1 gene therapy holds promise as translational treatment for Alzheimer’s disease, and that the concurrent administration of scFvs targeting early-conformational and late-phosphorylation changes does not result in any advantage in preclinical models of anti-tau immunotherapy, compared to targeting MC1-misfolded tau alone. It remains to be seen if testing other combinations of scFv and/or intrabodies directed to early and late phosphorylations, as well as scattering treatments over time sequentially could help maximize the effect of a dual treatment. We assume to have engaged intracellular tau, given the above discussion and the previous evidence of scFv intracellular localization. Future studies will be directed to ameliorate this system by further engineering our scFv, or nanobodies, to also express a transferrin receptor (TfR) recognition sequence, and improve blood brain barrier penetration.

## Materials and Methods

### ScFv-PHF1 design, characterization and sub-cloning into AAV1

As previously published for scFvMC1 (32), the light and heavy chain variable domains (V_H_ and V_L_) corresponding to the PHF1 antibody were sequenced using the MCLAB antibody service (San Francisco, CA). A 15-amino acid residue linker (Gly_4_Ser)_3_ was used to connect the V_H_ and V_L_ chains. A 5’-terminal signal peptide (SP) and 3’-terminal Myc and His6X tags were added. The scFvPHF1 purification procedure was performed as previously published (32). Briefly, scFvPHF1 was cloned into the mammalian expression vector pcDNA3.1 (Genewiz, South Plainfield, NJ). Upon transfection into HEK293T using Lipofectamine 2000 (Invitrogen, Carlsbad, CA), the scFv was released into the conditioned medium and affinity-purified using a Ni-Sepharose High Performance column (GE Healthcare, Port Washington, NY). The purified scFvPHF1 was analyzed on a Coomassie-stained SDS-PAGE gel to conform its molecular weight and purity. Tranfected cells were lysed using a solution of Tris-buffered saline (TBS), pH 7.4, containing 10 mM NaF, 1 mM Na_3_VO_4_ and 2 mM EGTA (plus complete Mini protease inhibitor cocktail, Roche), with 0.1% SDS added. The lysate underwent three 10-s cycles of sonication. Samples were run on 4-20% Criterion Tris-HCl gels (Bio-Rad Laboratories, Hercules, CA) and electrophoretically transferred to a nitrocellulose membrane (Thermo Fisher Scientific, Waltham, MA). Residual protein-binding sites were blocked by incubation with 5% non-fat milk in 1X TBST (1X TBS plus 0.1% Tween 20) for 1h at RT, followed by overnight incubation at 4°C with primary antibodies diluted in 20% SuperBlock buffer (Thermo Fisher Scientific, Waltham, MA) in 1X TBST. Anti-Myc-tag 9B11 (Cell Signaling, Danvers, MA) was diluted 1:1000. Appropriate isotype-specific HRP-conjugated secondary antibodies were diluted 1:10000 in 5% non-fat milk 1X TBST and added for 1 h at RT. Every step was followed by 3–4 washes in 1X TBST. Detection was performed using Pierce ECL Western Blotting Substrate (Thermo Fisher Scientific, Waltham, MA) and exposed to x-ray films. The parental antibody PHF1 and its recombinant version scFvPHF1 were compared in IHC on mouse brains (see details below). The antigen-binding specificity of scFvPHF1 was compared to scFvMC1 using an immunosorbent assay: 96-well plates were coated overnight with AD-derived PHF-tau (23, 58). Purified scFv-MC1 and scFvPHF1 were diluted in 5% non-fat milk and added to the wells in serial dilutions. After incubating for 1 h at RT, mouse anti-Myc-tag-HRP was added (1:1000 dilution) (Thermo Fisher Scientific, Waltham, MA). Bio-Rad HRP Substrate Kit (Bio-Rad laboratories, Hercules, CA) was used for detection and plates were read with an Infinite m200 plate reader (Tecan, San Jose, CA) at 415 nm (Supplementary Figure 1C). AAV packaging and purification services were provided by Vector Biolab (Malvern, PA): scFv-PHF1 was sub-cloned into the adeno-associated viral vector serotype 1 (AAV1) under the control of the synthetic strong CAG (CMV-chicken beta actin-rabbit beta globin) promoter. The Woodchuck hepatitis virus post-transcriptional regulatory element (WPRE) was added 5’ of the Myc and His6X tags to enhance transgene expression.

### Tau transgenic mice

All animal husbandry procedures performed were approved by the Institutional Animal Care and Use Committee. Homozygous P301S breeders were obtained from Dr. Michel Goedert (Cambridge, UK) (59). These mice, on pure C57BL/6 background, express 0N4R human tau carrying the P301S mutation under the control of the neuron-specific murine Thy-1 promoter. Homozygous P301S (both females and males) were used to assess tau clearance in the brain, motor and behavioral phenotype amelioration. A prevention paradigm assessed the treatment efficacy in the brain and the behavioral phenotype, in which injections were perfomed at 2 months of age, with sacrifice 4 months later. The following mice were included: (1) Sham group injected with eGFP (n = 17 mice, 8 females and 9 males), (2) scFvPHF1 monotherapy group (n = 14 mice, 7 females and 7 males), (3) scFvMC1 monotherapy group (n = 15 mice, 6 females and 9 males), and (4) Combination scFvMC1/scFvPHF1 group (n = 21 mice, 8 females and 13 males). A baseline cohort sacrificed at 2-month-old was also added to this study (n = 10, 5 females and 5 males). A small cohort of homozygous P301S mice was treated following the prevention paradigm described above; this group was used to pool the two hippocampi from each mouse and generate insoluble tau preparation (sham group: 6 mice, scFvMC1 monotherapy group: 7 mice). Mice were used to image ^18^F-FDG uptake using MicroPET: a) preliminary baseline study included 5.5-month-old C57BL/6 mice (n = 6, 4 females and 2 males) and P301S (n = 7, 4 females and 3 males); b) preliminary efficacy study included 5.5-month-old P301S mice injected with AAV1-CAG-eGFP (n = 7, 4 females and 3 males) or AAV1-CAG-scFvMC1 (n = 7, 4 females and 3 males).

### Intra-muscular (IM) injections

AAV1-CAG-scFvMC1, AAV1-CAG-scfvPHF1 and AAV1-CAG-eGFP (sham group) were each injected at a dose of 2×10^11^ GC per mouse. Each AAV was diluted in PBS to a final volume of 50 µL. Following a monotherapy paradigm a one-time IM injection was administrated in the right tibialis anterior muscle. For the combination therapy, AAV1-CAG-scFvMC1 was injected in the right tibialis anterior, while AAV1-CAG-scFvPHF1 was administered in the left tibialis anterior. All injections were performed under isoflurane anesthesia.

### Brain extracts and tissues preparation

Mice were sacrificed by isoflurane overdose, decapitated and processed as described previously (22). The brain was removed and divided at the midline, with half of the brain dissected for biochemical analysis. Cortex and hippocampus were homogenized separately using an appropriate volume of homogenizing buffer, consisting of Tris-buffered saline (TBS), pH 7.4, containing 10 mM NaF, 1 mM Na_3_VO_4_ and 2 mM EGTA, plus the complete Mini protease inhibitor cocktail (Roche). Brain homogenate aliquots were stored at −80°C and used for separate measurement of total protein concentration (DC Protein Assay, Bio-Rad Laboratories, Hercules, CA), soluble tau, and insoluble tau. Soluble tau was measured as heat-stable preparation (hsp) and was prepared by adding 5% ß-Mercaptoethanol and 200 mM NaCl to the brain homogenates. Samples were heated at 100°C for 10 min and then cooled at 4°C for 30 min. After centrifuging at 14,000g in a table-top microcentrifuge at 4°C for 10 min, supernatants were collected. To obtain insoluble tau preparations (INS), homogenates were thawed and centrifuged at maximum speed for 10 min at 4°C in a tabletop centrifuge. The collected supernatants were centrifuged at 200,000g for 30 min at 4°C. The resulting pellets were re-suspended in homogenizing buffer and centrifuged again at 200,000g for 30 min at 4°C. The final pellets were re-suspended in 1X sample buffer (5X sample buffer: Tris-buffered saline, pH 6.8, containing 4% SDS, 2% β-mercaptoethanol, 40% glycerol and 0.1% bromophenol blue) and heated at 100°C for 10 min to efficiently dissociate the insoluble tau fraction. Of note, right and left hippocampi from each mouse were pooled in order to extract insoluble tau from this region.

### Tau ELISA

Levels of total, phosphorylated and aggregated tau were assessed using the Low-tau ELISA (enzyme-linked immunosorbent assay) protocol previously published (45, 60). 96-well plates were coated for 48 h at 4°C with specific purified monoclonal tau antibodies (DA31, CP13, PHF1, RZ3, DA9) at a concentration of 6µg/mL. After washing, plates were blocked for 1 h at RT using StartingBlock buffer (Thermo Fisher Scientific, Waltham, MA). Brain samples and standards were diluted in 20% SuperBlock buffer (Thermo Fisher Scientific, Waltham, MA) in 1X TBS and loaded onto the plates. The total tau detection antibody DA9-HRP, diluted 1:50 in 20% SuperBlock in 1X TBS, was added to the samples and tapped to combine. For the aggregated/oligomeric tau ELISA using DA9 both as capture and detection antibody, the DA9-HRP was added sequentially the following day. To visualize the ELISA, 1-Step ULTRA TMB-ELISA (Thermo Fisher Scientific, Waltham, MA) was added for 30 min at RT, followed by addition of 2N H_2_SO_4_ to stop the reaction. Plates were read with Infinite m200 plate reader (Tecan, San Jose, CA) at 450nm.

### Immunohistochemistry

Tau staining was performed according to standardized protocols (22, 32). After decapitation, one hemisphere was fixed overnight in 4% paraformaldehyde at 4°C. Using a vibratome, serial sections were cut and stained as free-floating in 24-well plates using a panel of tau antibodies. To control for potential scFvMC1 antigen-masking, antigen retrieval was performed using 1X Dako Target Retrieval solution (Agilent Dako, Santa Clara, CA, USA) in distilled water/0.5% Triton, at 95–99°C for 5 min. After washing, endogenous peroxidases were quenched with 3% H_2_O_2_/0.25% Triton X-100/1X TBS for 30 min. Non-specific binding was blocked with 5% non-fat milk-1X TBS for 1 h at RT. Primary anti-tau antibodies were diluted, MC1 (1:500) and PHF1 (1:5000), in 5% non-fat milk-1X TBS, and incubated overnight at 4°C with shaking. Biotin-conjugated secondary antibodies (SouthernBiotech, Birmingham, AL) directed against the specific isotypes were diluted 1:1000 in 20% SuperBlock, incubated for 2 h at RT, and followed by Streptavidin-HRP (SouthernBiotech, Birmingham, AL) incubation for 1h. Staining was visualized using 3,3’-Diaminobenzidine (Sigma-Aldrich, St. Louis, MO). For immuno-staining using purified scFv-PHF1 (1:1000), an anti-Myc-Tag mouse mAb (clone 9B11) (Cell Signaling Danvers, MA) was used, followed by the specific biotin-conjugated secondary antibody (IgG2a). All other steps in the protocol remained the same. Images were acquired using Olympus BH-2 bright field microscope, analyzed and processed using ImageJ/Fiji software (NIH). Semi-quantification was performed on frontal cortex and dentate gyrus using the measure particles tool, working with 8-bit images and adjusting the threshold. A threshold of 0–255 and particle size of 150-infinity was applied to all cortical and dentate gyrus images for consistency.

### Hind-limbs clasping scale

Hind-limb reflex is a functional motor test used to quantify deficits in corticospinal function. Limb clasping is observed in several transgenic tau mouse models (59, 61–64) and recapitulates some of the functional motor deficits observed in late-stage AD patients. Suspension of mice by the tail normally triggers an escape response (score = 0). Deficits in the ability to display the described extension are scored based on their severity on a clasping scale (from 0 to 5). When suspended by the tail, an animal with a partial limb extension reflex will start reducing its hind-limbs’ extension gradually, progressing from a score 1 to 5 where a complete absence of extension reflex in both hind-limbs is observed and paresis occurs shortly thereafter. Mice were monitored weekly for clasping. At sacrifice, mice were scored to assess afficacy of scFv treatments.

### Open-field task

All mice used for behavioral assays were housed under a reversed dark (9:00–21:00) and light (21:00–9:00) cycle, with ad libitum access to food and water. We assessed 4 different treatment groups: Sham, scFvPHF1, scFvMC1 and combination scFvMC1/PHF1. Mice were studied at 16 and 23 weeks of age. All manipulations were conducted during the dark phase, at least 1 h after turning lights off, and male and female mice were assessed on different days. Prior to behavioral assessments, mice were handled three times, for 15 min each, on separate days. The open-field task (65) was used to examine locomotor activity, and anxiety-like behavior by placing the mice in the center of a square arena (40 × 40 cm^2^) with gray walls (35 cm high) and allowing them to freely explore the chamber during a 10-min session. The sessions were recorded with a centrally placed video camera directly above the arena which fed the signal to the tracking software (EthoVision XT, version 15, Noldus, Attleboro, MA, USA) used for automated analysis of animal behaviors including distance traveled, velocity, time spent moving, mobility (a measurement of how much an animal moves around, calculated by analyzing the change in its body area between successive frames of a video), time spent in the center of the arena (18.9 × 18.9 cm^2^) and the four corners of the arena. Experiments were conducted and analyzed according to randomly assigned cage numbers which did not indicate the scFv exposure.

### Rotarod task

This task measured motor coordination and balance (65)by placing 16-week-old mice on a five-lane rotating drum (ENV-574M, Med Associates Inc, St. George, VT, USA), which accelerated from 4 to 40 rpm over a course of 5 min. The riding time until each mouse fell off the drum was recorded. The test was repeated 5 times for each mouse during 5 consecutive days (one trial per day). The room was illuminated with low-level orange lights.

### 18F-FDG-PET and Imaging reconstruction

Imaging was performed using the Siemens Inveon® positron emission tomograph (MicroPET) system. The radiotracer ^18^F-FDG is synthesized onsite at the PET imaging facility at the Feinstein Institutes and delivered directly into the microPET suite. All animals were fasted overnight (8-12 hours) prior to PET imaging to stabilize blood glucose levels. On imaging day, mice were acclimated for one hour upon arrival to the imaging suite. Two mice were simultaneously anesthetized with induction at 2-2.5% isoflurane mixed in oxygen and maintained at 1.5-2.0% with custom nose cone allowing for head-to-head configuration. After confirming depth of anesthesia, approximately 0.5 mCi of ^18^F-FDG (in 0.3ml) was injected i.p. with 35-40 mins allowed for uptake of the tracer followed by a 10 min static scan.

Brain images were acquired as 600 second static emission with transmission attenuation correction (given that the animals were immobile on the gantry over the course of imaging acquisition). Reconstruction was completed using Inveon Acquisition workflow (Siemens AW 1.5) with three-dimensional ordered subsets-expectation maximization-maximum *a posteriori* recontruction, (OSEM3D-MAP, 20 iterations) at a target resolution of 1.5mm. After reconstruction, the raw images (final matrix of 128 × 128 × 159mm with pixel size of 0.78 × 0.78 × 0.8 mm) were pre-processed using Pixel-Wise Modeling software (PMOD 4.0) in the following way: raw images were bounding box aligned to an anatomical template, skull stripped, and dose and weight corrected. ^18^FDG scans from each animal were registered to an ^18^FDG template and then to a C57/BL6J MRI template (66), both of which were aligned within Paxinos and Franklin anatomical space. The template-aligned ^18^FDG scans were then applied to C57/BL6J MRI template for rigid transformation using statistical parametric mapping (SPM5) with SPMmouse toolbox within MATLAB. Images were smoothed with an isotropic Gaussian kernel FWHM (full width at half maximum) 0.2 mm at all directions. Group statistical differences were considered significant at a voxel-level threshold of *P* < 0.05 and a cluster cutoff between 50-400 voxels for increases and decreases.

To identify brain regions in anatomical space with significant differences between mice groups (C57BL/6 vs P301S, Sham vs scFvMC1), we performed whole brain voxel-wise search, two sample pairings using SPMmouse software within MATLAB computing environment (The Wellcome Centre for Human Neuroimaging, UCL Queen Square Institute of Neurology, London, UK, https://www.fil.ion.ucl.ac.uk/spm/ext/#SPMMouse). SPM parameters for alignment and anatomical registration to mean were as follows: quality: 0.9, separation: 0.33, smoothing 0.2 in all directions, no wrap, maksed using MRI mouse template in Paxinos and Franklin coordinates. Statistically significant cluster extraction using FSLeyes (University of Oxford, Oxford, UK, https://fsl.fmrib.ox.ac.uk/fsl/docs/#/utilities/fsleyes)

### Statistical analysis

Quantitative and semi-quantitative data (Tau ELISAs, IHC) were analyzed using the dedicated software GraphPad Prism V.10 (GraphPad software Inc., CA). Ordinary One-way ANOVA with Tukey’s multiple comparison test was performed when the parametric assumption of normality (D’Agostino-Pearson omnibus test) was accomplished. When not, non parametric Kruskal-Wallis test with Dunn’s correction was run instead. For behavioral assays (rotarod, open-field task), we used two-way ANOVA with repeated measures (RMANOVA), with treatment and individual trials as factors, followed by Tukey post-hoc test. Statistical significance was set at *P* < 0.05. Error bars epresent the standard error of the mean (SEM). Two tailed unpaired t-test with Welch’s correction was used for ^18^F-FDG-PET analysis.

## Data Availability statement

The datasets used and/or analyzed during the current study are available from the corresponding authors upon reasonable request.

## Acknowledgements

This work was supported by funding from the National Institute of Health, National institute on Aging (NIA) grant 5R01AG061381 (to CD). Further funding was provided by the National Institute of Health (NIH) grant 5P01AI102852 (to PH) and NIH grant 5P01AI073693 (to PH) as well as the Department of Defense (DOD) impact award W81XWH1910759 (to PH). The authors would like to recognize the late Dr Peter Davies for the anti-tau hybridoma available at the Feinstein Institutes for Medical Research. We thank the Feinstein Institute’s Cyclotron/Radiochemistry Department, Dr Tom Chaly, Prasad Majji, Matthew Hellman, and the CCP for assistance with cannulation from Francisco Colon and Meaghan Bragg.

## Authors’ contributions

SK performed experiments, analyzed data and helped preparing the manuscript; JC performed experiments, analyzed data; VV performed experiments and edited the manuscript; JC perfomed microPET, analysed data and helped preparing the manuscript; LM helped analyse microPET data; PH perfomed rotarod and open field tasks, interpreted data, prepared figures and helped preparing the manuscript; PM interpreted experiments and edited the manuscript; LG interpreted experiments and prepared the manuscript; CD designed the overall project, performed experiments, analyzed data, interpreted results and prepared the manuscript. All authors read and approved the final manuscript.

## Declaration of interests statement

The authors declare no conflict of interest.

---

**Figure S1.**
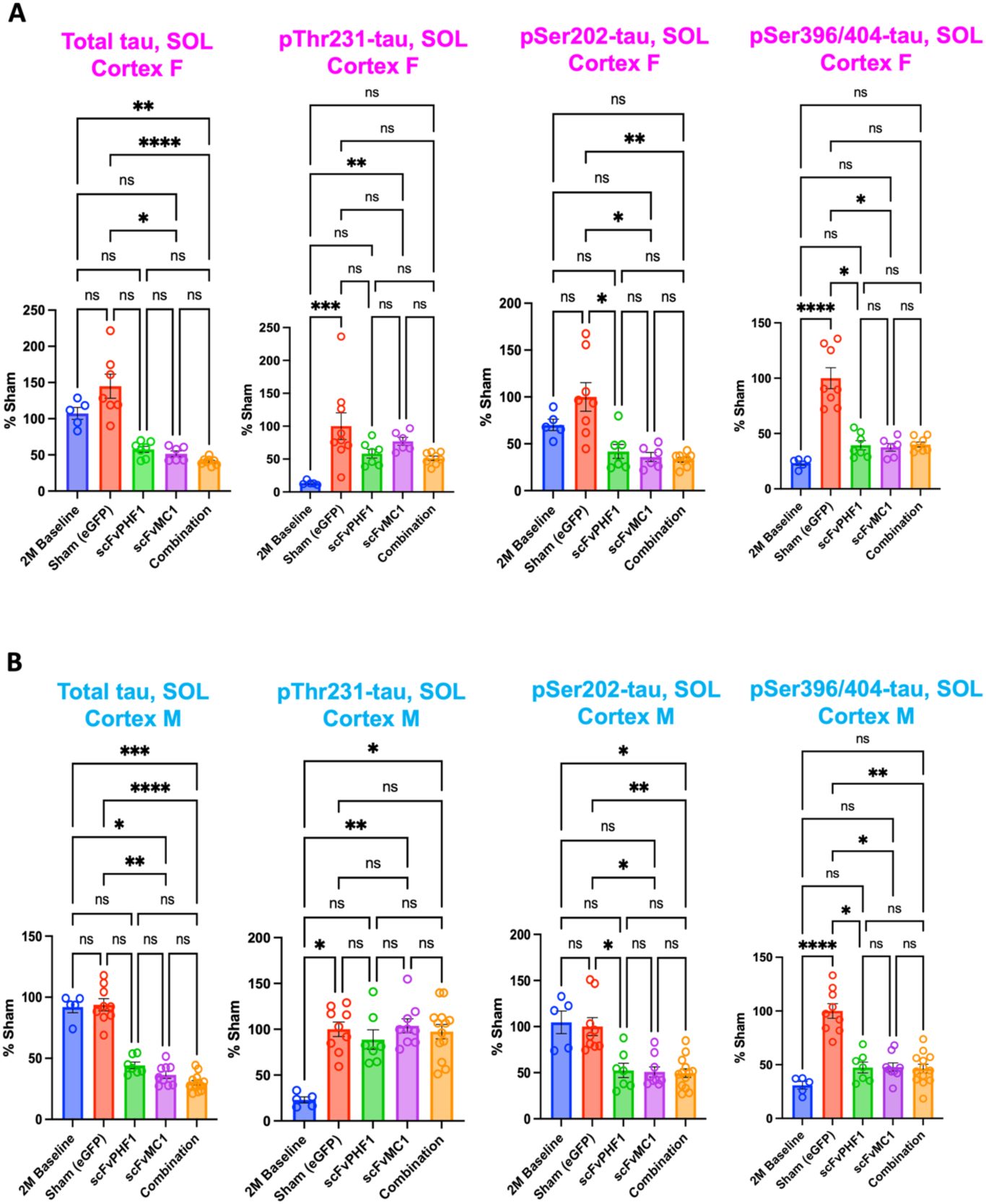
scFv-MC1 reduces soluble tau in the cortex without major sex differences, related to Figure 2. **(A)** ELISA quantifications of soluble (SOL) total tau, pThr231-tau, pSer202-tau, and pSer396/− 404-tau in females: baseline (n = 5), sham-eGFP (n = 9), scFv-PHF1 (n = 7), scFv-MC1 (n = 6), and combination (n = 8). **(B)** ELISA quantifications of soluble (SOL) total tau, pThr231-tau, pSer202-tau, and pSer396/-404-tau in males: baseline (n = 5), sham-eGFP (n = 9), scFv-PHF1 (n = 7), scFv-MC1 (n = 9), and combination (n = 13). Data were normalized to the percent sham-eGFP group and represent mean ± SEM. Statistical analysis was performed by parametric Kruskal-Wallis test. * *P* < 0.05; ** *P* < 0.01; *** *P* < 0.001; **** *P* < 0.0001.

**Figure S2.**
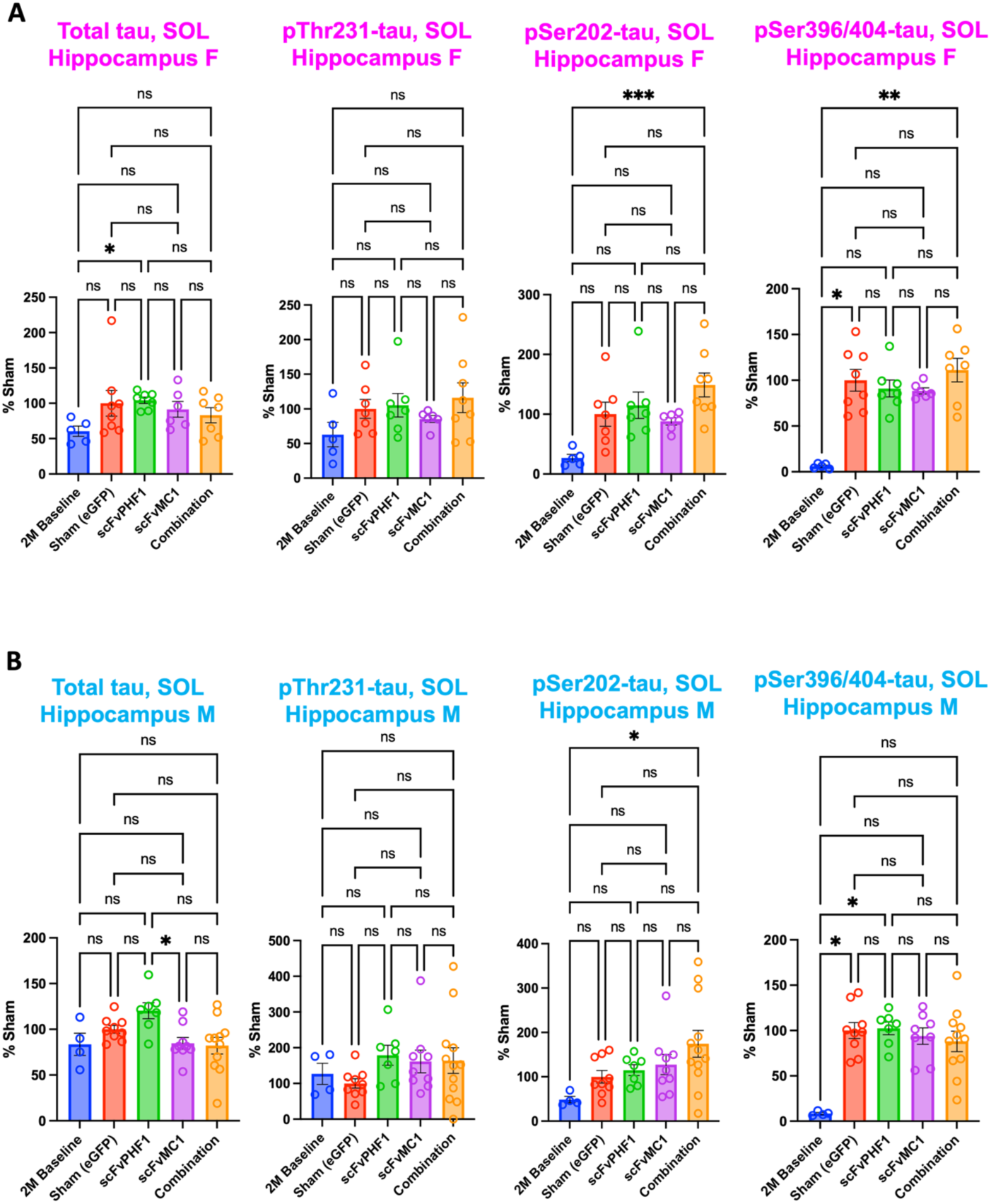
Hippocampal soluble tau remains relatively unchanged across sexes, related to Figure 3. **(A)** ELISA quantifications of soluble (SOL) total tau, pThr231-tau, pSer202-tau, and pSer396/-404-tau in females: baseline (n = 5), sham-eGFP (n = 8), scFv-PHF1 (n = 7), scFv-MC1 (n = 6), and combination (n = 8). **(B)** ELISA quantifications of soluble (SOL) male total tau, pThr231-tau, pSer202-tau, and pSer396/− 404-tau in males: baseline (n = 5), sham-eGFP (n = 9), scFv-PHF1 (n = 7), scFv-MC1 (n = 9), and combination (n = 12). Data were normalized to the percent sham-eGFP group and represent mean ± SEM. Statistical analysis was performed by parametric Kruskal-Wallis test. * *P* < 0.05; ** *P* < 0.01; *** *P* < 0.001.

**Fig S3.**
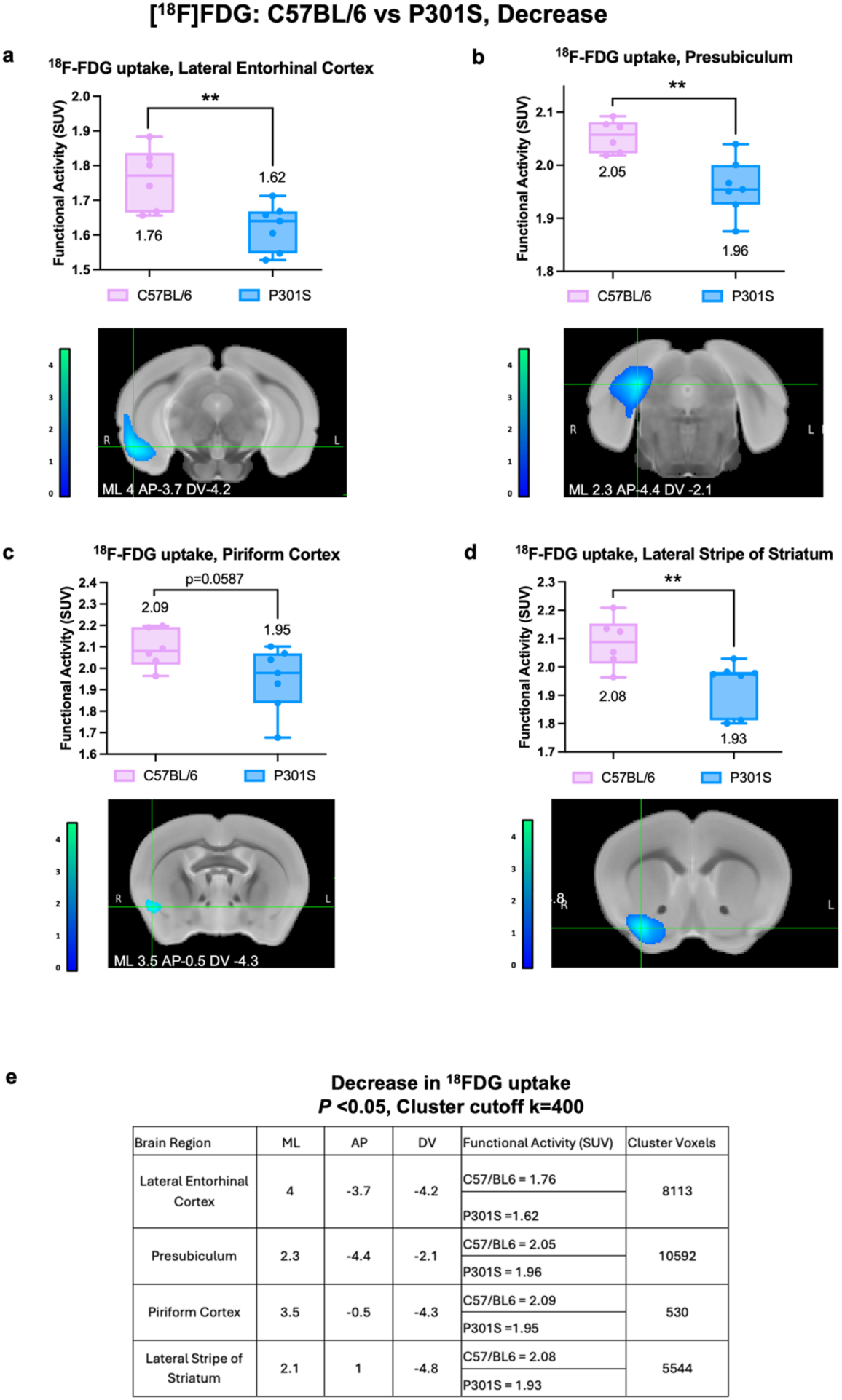
Brain regions with decreased ^18^F-FDG uptake in C57BL/6 vs P301S mice, related to Figure 7. Coronal view of statistically significant decrease C57 BL/6 mice (n=6) vs. P301S (n=7) in relative ^18^F-FDG uptake at P < 0.05. Regions with lower metabolic activity includes lateral entorhinal cortex **(a)**, presubiculum **(b)**, piriform cortex **(c)**, and lateral stripe of striatum **(d)**. Statistically significant clusters were extracted using FSLeyes and overlaid onto coronal view mouse brain MRI template, Paxinos & Franklin anatomical coordinates shown. Color bar representation of ^18^F-FDG decrease threshold value. Data are expressed as the mean + SEM. * *P* < 0.05, ** *P* < 0.01, *** *P* < 0.001, two tailed unpaired t-test with Welch’s correction. **(e)** Brain regions in Paxinos and Franklin coordinates for significant clusters of ^18^FDG activation: table includes stereotactic coordinates: ML (mediolateral), AP (anterior posterior), DV (dorsoventral), functional activity average in C57/BL6 and P301S, and cluster voxels. SPM derived cluster value (standardized uptake values, SUV), *P* <0.05, Cluster cutoff k=400.

**Figure S4.**
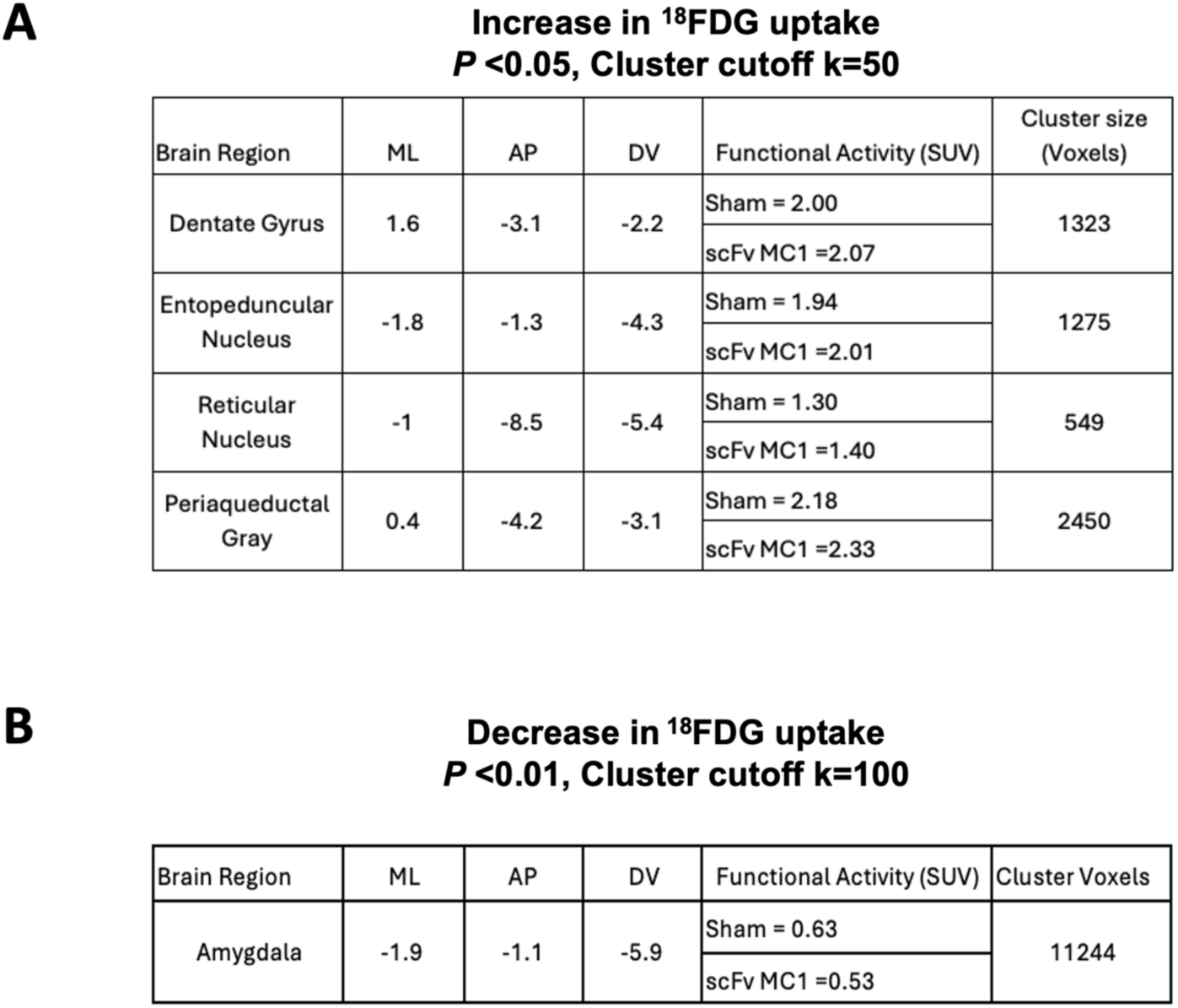
^18^F-FDG uptake: brain regions, functional activity average and cluster voxels in P301S treated and sham-eGFP mice, related to Figure 7. **(A)** Brain regions in Paxinos and Franklin coordinates for significant clusters of ^18^FDG activation. Increase in ^18^FDG uptake in scFvMC1 treated mice. SPM group comparison in Sham (n=7) vs scFv MC1 (n=7), SPM derived cluster value (standardized uptake values, SUV), p<0.05 with voxel cutoff at k=50. **(B)** Brain regions in Paxinos and Franklin coordinates for significant clusters of ^18^FDG activation. Decrease in ^18^FDG uptake in scFvMC1 mice. SPM group comparison in Sham, (n=7) vs scFv MC1 (n=7), SPM derived cluster value (standardized uptake values, SUV), p<0.01, Cluster cutoff k=100. ML (mediolateral), AP (anterior posterior), DV (dorsoventral).

## Notes

### Competing Interest Statement

The authors have declared no competing interest.

